# Host carbon dioxide concentration is an independent stress for *Cryptococcus neoformans* that affects virulence and antifungal susceptibility

**DOI:** 10.1101/659789

**Authors:** Damian J. Krysan, Bing Zhai, Sarah R. Beattie, Kara M. Misel, Melanie Wellington, Xiaorong Lin

**Affiliations:** Department of Pediatrics, Carver College of Medicine, University of Iowa, Iowa City, Iowa 52242; Microbiology/Immunology, Carver College of Medicine, University of Iowa, Iowa City, Iowa 52242; Department of Biology, Texas A&M University, College Station, TX 77843; Program in Molecular Medicine, Carver College of Medicine, University of Iowa, Iowa City, IA 52242; Department of Microbiology, University of Georgia, Athens, GA, 30602

**Author notes:** Corresponding Authors: Damian J. Krysan, 2040 Med Labs 25 S. Grand Avenue, Department of Pediatrics and Microbiology/Immunology, Carver College of Medicine; University of Iowa, Iowa City, IA 52242, Xiaorong Lin, 208 Biological Sciences Building, Department of Microbiology, University of Georgia 120 Cedar Street, Athens GA 30602.

## Abstract

The ability of *Cryptococcus neoformans* to cause disease in humans varies significantly among strains with highly related genotypes. In general, environmental isolates of pathogenic species such as *C. neoformans* var. *grubii* have reduced virulence relative to clinical isolates, despite having no differences in the expression of the canonical virulence traits (high temperature growth, melanization and capsule formation). In this observation, we report that environmental isolates of *C. neoformans* tolerate host CO_2_ concentrations poorly compared to clinical isolates and that CO_2_ tolerance correlates well with the ability of the isolates to cause disease in mammals. Initial experiments also suggest that CO_2_ tolerance is particularly important for dissemination of *C. neoformans* from the lung to the brain. Furthermore, CO_2_ concentrations affect the susceptibility of both clinical and environmental *C. neoformans* isolates to the azole class of antifungal drugs, suggesting that antifungal testing in the presence of CO_2_ may improve the correlation between in vitro azole activity and patient outcome.

**Importance:** A number of studies comparing either patient outcomes or model system virulence across large collections of *Cryptococcus* isolates have found significant heterogeneity in virulence even among strains with highly related genotypes. Because this heterogeneity cannot be explained by variations in the three well-characterized virulence traits (growth at host body temperature; melanization; and polysaccharide capsule formation), it has been widely proposed that additional *C. neoformans* virulence traits must exist. *C. neoformans* natural niche is in the environment where the carbon dioxide concentration is very low (∼0.04%); in contrast, mammalian host tissue carbon dioxide concentrations are 125-fold higher (5%). We have found that the ability to grow in the presence of 5% carbon dioxide distinguishes low virulence strains from high virulence strains, even those with a similar genotype. Our findings suggest that carbon dioxide tolerance is a previously un-recognized virulence trait for *C. neoformans*.

## Introduction

*Cryptococcus neoformans* is one of the most important human fungal pathogens and causes meningoencephalitis (CME). Recent estimates indicate that 223,000 new cases of CME occur each year with an annual mortality of 181,000 (1); the majority of CME disease affects people infected with HIV (2). *Cryptococcus* species are environmental yeasts that occupy a variety of niches and, therefore, *Cryptococcus* must transition from an environmental niche to the mammalian host to cause disease (3). Most strains isolated from the environment are much less virulent in animal models when compared to strains isolated from human patients. For example, Litvintseva and Mitchell (4) found that only one out of ten environmental strains caused mortality in a murine model of cryptococcosis by 60 days while 5/7 clinical strains caused lethal infection by 40 days. In vitro, all of the strains grew at 37°C and generated comparable levels of capsule and melanin (4).

Recently, Mukamera et al. systematically characterized a set of genetically similar strains associated with highly variable patient outcomes (5). The virulence trends observed in patients were recapitulated in the inhalational murine model, providing important validation of the model as predicative of clinical virulence. Like Litvintseva and Mitchell (4), extensive phenotyping of the strains did not reveal an in vitro phenotype that correlated with virulence (5). These data strongly indicate that uncharacterized virulence properties beyond the “big three” of host body temperature tolerance, melanization, and capsule formation plays an important role in determining the virulence potential of a given cryptococcal strain (6).

We hypothesized that the host environment may contain additional stresses under which clinical/pathogenic isolates are fitter than environmental/non-pathogenic isolates. One dramatic difference between terrestrial and host environments is the concentration of carbon dioxide (CO_2_): ambient air is ∼0.04% CO_2_ while the CO_2_ concentration in mammalian tissues is 125-fold higher (5%). In ambient air, Cryptococcus, like many yeasts expresses carbonic anhydrase (Can2) which catalyzes the generation of essential HCO_3_^−^ under low CO_2_ concentrations. At 5% CO_2_ in the host, *CAN2* expression is repressed presumably due to sufficient levels of HCO_3_^−^ produced by dissolved CO_2_. Consequently, *CAN2* is dispensable for growth at 5% CO_2_ and virulence in a murine host (10). Host CO_2_ concentrations, however, have a profound effect on *C. neoformans* biology because they induce capsule formation (7). Indeed, our hypothesis that host concentrations of CO_2_ are a significant stress to *C. neoformans* is supported by the fact that Granger et al. noted that the growth of cultures slowed considerably after being shifted to host CO_2_ concentrations to induce capsule formation (8). In addition, *C. neoformans* strains lacking calcineurin, a key stress response regulator, are hypersensitive to CO_2_ (9). Finally, Bahn et al. found that elevated concentrations of CO_2_ inhibit *C. neoformans* mating (10). Taken together, tolerance of host CO_2_ concentrations seemed to us a potentially important independent trait of *C. neoformans* strains that cause disease in mammals.

## Results and Discussion

To test the hypothesis that CO_2_ tolerance may play a role in distinguishing between clinical and environmental strains of *C. neoformans*, we examined the CO_2_ fitness of a set of 12 strains that had been previously characterized in the mouse pulmonary infection model by Litvintseva and Mitchell (4). The clinical and reference strain H99 as well as three other clinical strains grew similarly in the presence and absence of CO_2_ on RPMI media buffered to pH 7 with MOPS while all but one of the environmental strains displayed a growth defect in 5% CO_2_ (Fig. 1A). The serotype D reference strain JEC21 is also CO_2_ sensitive relative to H99 and the clinical isolates. Of 10 additional clinical strains isolated from patients at Duke University (gift of John Perfect), 8 had similar growth at ambient and host CO_2_ concentrations (Fig. S1). Of the 12 strains that were examined by Litvintseva and Mitchell in animal models, only those that were CO_2_-tolerant caused disease by 60 days. The only environmental strain to be CO_2_ tolerant was also the only one to cause disease.

**Figure 1.**
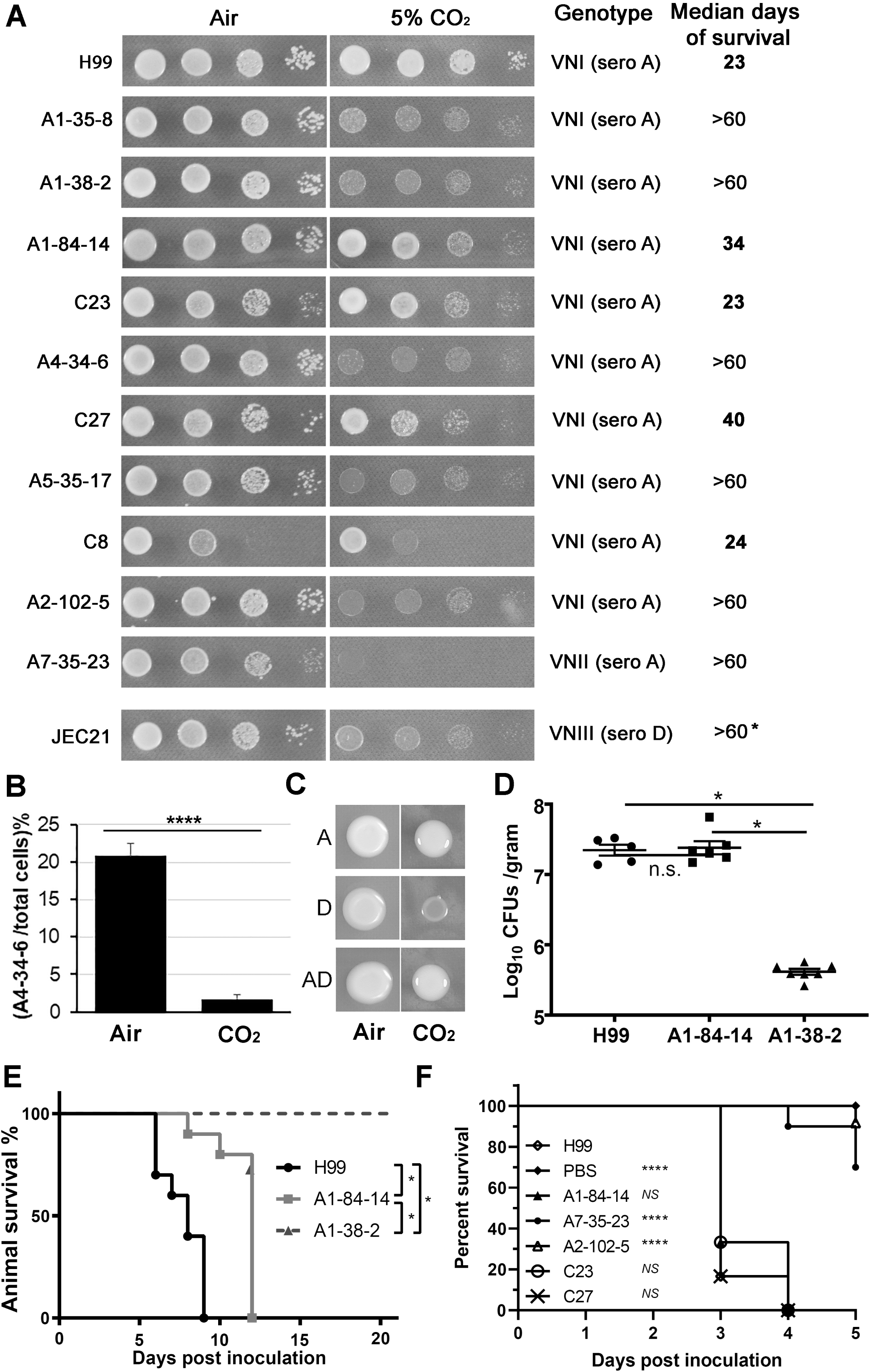
*In vitro* CO_2_-tolerance correlates with cryptococcal virulence in insect and mammalian hosts. (**A**) Clinical and environmental *C. neoformans* isolates were cultured on RPMI medium buffered to pH 7 with MOPS at 37°C in ambient air (∼0.04% CO_2_) or in 5% CO_2_. The information about the genotype of the strains and the median survival days of mice infected by these strains intranasally were obtained from a previous study (4). (**B**) Competition assay of isolate A4-34-6 and mCherry labeled H99 cultured on RPMI medium buffered to pH 7 with 165 mM MOPS in ambient air or in 5% CO_2_. *P*< 0.00001, Student’s t test. (**C**) Cell suspensions of H99 (A), JEC21 (D), and XL1462 (AD hybrid) at equal concentration were spotted onto RPMI medium and incubated at 37°C in ambient air or in 5% CO_2_. (**D &E**) CD-1 mice were inoculated with H99, CO_2_ tolerant environmental strain A1-84-14, and CO_2_ sensitive strain A1-38-2 (5-7 animals per group) by tail vein injection. Five animals were sacrificed at 5 days for brain fungal burden and ten animals per group were monitored for morbidity until day 21 (**D**); * indicates statistically significant difference between indicated group by Student’s t test of log-transformed data. Survival curves were analyzed by Kaplan-Meier/Log-rank test (**E**). (**F**) The great wax moth *Galleria mellonella* larvae in the final instar larval stage (10-15 per group) were injected with the indicated clinical or the environmental isolates *via* the last left proleg. Survival rate of the infected larvae over days post inoculation is shown. *NS*: non-significant; \*\*\*\**P*<0.0001 compared to H99 by Log-rank test.

To obtain a more quantitative measure of the in vitro fitness advantage of a CO_2_ tolerant strain over a CO_2_-sensitive strain, we carried out a competition experiment in which mCherry-labeled H99 strain and unlabeled, environmental strain A4-34-6 were co-cultured as a 1:1 mixture in ambient air or 5% CO_2_; the ratio of H99 to A4-34-6 was determined by microscopy. A4-34-6 has a 5-fold fitness defect relative to H99 in ambient air and that defect is increased to 50-fold in 5% CO_2_ (Fig. 1B). To determine if CO_2_ tolerance is recessive or dominant, the previously generated diploid AD hybrid was compared to its parental strains serotype A H99 and serotype D JEC21 (11). The diploid AD hybrid is CO_2_ tolerant indicating that the trait is dominant (Fig. 1C). Consistent with other yeasts (12), CO_2_ is fungistatic and dose dependent (data not shown). Because CO_2_/HCO_3_ concentrations^−^ are substrates in reactions of central carbon metabolism (12), we wondered if increasing the glucose concentration of RPMI from 0.2 to 2% would affect CO_2_ sensitivity; however, it had no effect on the growth of the sensitive strains (Fig. S1).

As reported by Litvintseva and Mitchell (4), all clinical strains caused lethal infections in mice within 40 days while the only environmental strain to cause a lethal infection within 60 days was the CO_2_ tolerant strain, A1-84-14 (Fig. 1A). To determine if the virulence differences between the CO_2_ tolerant and CO_2_ sensitive strains were dependent on the infection model, the environmental CO_2_ tolerant and CO_2_ sensitive strains were compared in both the intravenous model of disseminated murine cryptococcosis and in the *Galleria mellonella* model. Five days post-infection (Fig. 1D), the fungal brain burden was 2 log_10_ CFU/g lower in the CO_2_ sensitive environmental strain relative to the CO_2_ tolerant environmental strain, while the CO_2_ tolerant environmental strain was similar to the reference strain H99. Consistent with the pulmonary infection model data, the median survival of mice infected with the CO_2_ tolerant environmental strain is modestly longer than the highly virulent H99 reference strain (Fig. 1E). Mice infected with the CO_2_ sensitive strain, in contrast, were asymptomatic for an additional week. The fungal burden for the CO_2_ sensitive strain infected mice had increased 1.7 log_10_ over the 16 days (Fig. S3), indicating that environmental strain replicated very slowly in the brain. Finally, *G. mellonella* larvae infected with CO_2_ sensitive strains showed prolonged survival relative to CO_2_ tolerant strains (Fig. 1F). Taken together, these data strongly support a correlation between CO_2_ tolerance and virulence in multiple models of cryptococcal infection.

CO_2_ levels affect the composition and function of cellular membranes in a variety of biological systems which is proposed to be one possible mechanism of CO_2_-mediated, growth inhibition (12, 13). The two most important anti-cryptococcal drugs, amphotericin B and fluconazole, affect membrane-related processes (14) and, thus, we hypothesized that CO_2_ may modulate antifungal susceptibility. The minimum inhibitory concentration (MIC) was determined using E-test strips on solid agar RPMI medium buffered to pH 7 with 165 mM MOPS. The MIC for amphotericin B was not affected by CO_2_ in any of the strains tested (≤ 2-fold change between ambient air and 5% CO_2_, Fig. 2A and Fig. S4B). In contrast, the fluconazole MIC decreased approximately 8 to 10-fold (4 µg/mL to 0.5 µg/mL) for H99, CO_2_-tolerant clinical isolates (C23 and C27) and environmental (A1-84-14) isolates in the presence of 5% CO_2_ (Fig. 2B and Fig. S4B). The MIC of both itraconazole (Fig. 2B) and voriconazole (Fig. S4B) is decreased at 5% CO_2_, indicating that the effect is not limited to fluconazole. 5% CO_2_ has no effect on the MIC of fluconazole against the *C. albicans* reference strain SC5314 (2 µg/mL). The effect of CO_2_ is also not specific to ergosterol biosynthesis inhibitors in that the sphingolipid biosynthesis inhibitor myriocin is also more active at host CO_2_ levels than ambient air (Fig. 2B).

**Figure 2.**
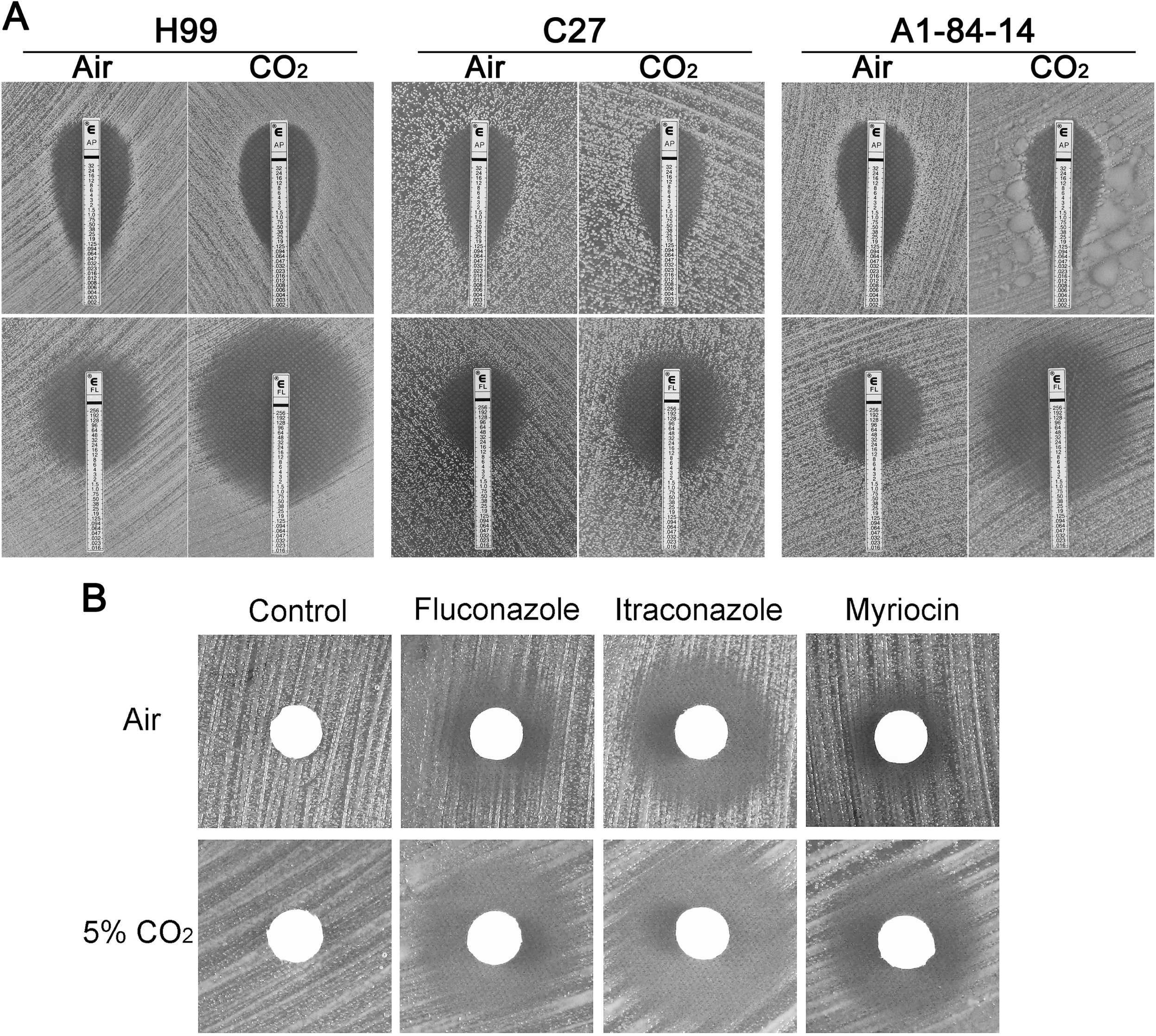
CO_2_ affects cryptococcal susceptibility to antifungals. (**A**) The CO_2_ tolerant cryptococcal isolates, including the reference and clinical isolate H99, the clinical isolate C27, and the environmental isolate A1-84-14, are more susceptible to fluconazole in 5% CO_2_ relative to ambient air but amphotericin B susceptibility is unaffected. Suspensions of *C. neoformans* isolates were spread onto RPMI agar medium. E-test strips with fluconazole (FL) or amphotericin B (AP) were placed on top of the air-dried yeast lawn. The cells were then incubated at 37°C in ambient air or in 5% CO_2_. (**B**) H99 is more susceptible to fluconazole, itraconazole, and myriocin in 5% CO_2_ by disk diffusion assay. H99 cells were spread onto RPMI agar medium. Disks containing fluconazole (4 μg), itraconazole (4 μg), myriocin (0.8 μg), or DMSO (control) were air dried and placed on top of the yeast lawn. Cells were incubated at 37°C for 2 days in ambient air or in 5% CO_2_. The size of the halo surrounding the disks correlates with cryptococcal susceptibility to the drugs.

The physiological mechanism underlying the differential sensitivity of Cryptococcus strains to host levels of CO_2_ awaits further study. A variety of potential mechanisms for the fungistatic and bacteriostatic effect of CO_2_ have been proposed including altered membrane fluidity and inhibition of biosynthetic reactions involving CO_2_ (12). Despite these outstanding mechanistic questions, the effects of host concentrations of CO_2_ on *C. neoformans* virulence and antifungal susceptibility have two important implications. First, CO_2_ tolerance appears to be an independent virulence feature of pathogenic strains of *C. neoformans* that correlates with variations in mammalian virulence. It will be interesting to expand this analysis to determine if CO_2_ tolerance contributes to the variation in patient outcome (5). Based on fungal burden data reported by Litvintseva and Mitchell (4), the CO_2_ sensitive strains cause a significantly lower lung burden than CO_2_ tolerant strains while the brain burden for CO_2_ sensitive strains is almost undetectable at post-infection day 60. Although the CO_2_ sensitive strain has reduced virulence in the intravenous model, which rapidly establishes CNS infection, it is able to replicate within the brain. Taken together, these preliminary and previously published data (4) indicate that CO_2_ tolerance may play a more important role in dissemination from the lung than in replication within the brain.

Second, the profound effect of host CO_2_ concentrations on in vitro azole activity identifies a potential limitation of current antifungal drug susceptibility testing conditions in predicting the outcomes of patients treated with azoles. Indeed, the lack of correlation between MICs generated by standardized antifungal susceptibility testing assays and clinical outcomes for cryptococcal infections has been well described and a number of potential explanations for this discrepancy have been proposed (15). Our data suggest that CO_2_ is likely to be an important factor in drug susceptibility. Although the mechanism of this effect will require additional investigation, it is unlikely that high CO_2_ concentrations directly reduce ergosterol levels because strains with reduced ergosterol content typically have reduced susceptibility to amphotericin B, a drug that binds directly to ergosterol. Regardless of the mechanism, the use of host CO_2_ concentrations may represent a simple adjustment that could improve the correlation between MIC and clinical outcome.

## Materials and Methods

### Strains and growth conditions

Media was prepared using standard recipes (16). Reference strains H99 and JEC21 were from stocks in the Krysan and Lin labs. Environmental and clinical strains were generous gifts from Anastasia Litvintseva, Tom Mitchell and John Perfect. Strains were stored in −80°C in 15% glycerol. Freshly streaked out yeast cells were grown on yeast peptone 2% dextrose (YPD) medium at 30°C. For spotting assays, the cells were washed, adjusted to the same cell density and serially diluted. Serial dilutions (3 µL) were then spotted onto RPMI without glutamine (buffered to pH 7 with 165 mM MOPS) agar plates and incubated at 37°C in ambient air or at 5% CO_2_. For the glucose supplementation assay, 2% glucose was added to RPMI medium buffered to pH 7 with 165 mM MOPS. H99 was labelled with mCherry by integrating plasmid pH3mCHSH2 (Addgene) into the so-called Safe-Haven 2 (SH2) locus described by Upadhya et al. (17). Transiently expressed Cas9 (18) was used to generate a double strand break in the SH2 region and pH3mCHSH2, linearized with ApaI, was used as the repair construct. After selection on neomycin containing plates, colonies that remained NEO**^+^** and fluorescent were isolated.

### Competition assay

mCherry-labeled H99 and A4-34-6 were grown overnight in liquid YPD at 30°C. The stationary phase cells were adjusted to identical cell density and a combined inoculum with equal numbers of cells was spotted on RPMI plates as described above. After 2 days in either ambient air or 5% CO_2_, samples of cells from different regions of the colony were collected and imaged in bright field (total cell number) and red channel using a Nikon epi-fluorescence microscope with a Cool Snap HQ2 camera, and Nikon Elements image acquisition and analysis software. The ratio of non-fluorescent to total cells was used to generate a competitive index (>100 cells counted for each data point). The reported data are from three biological replicates with three technical replicates.

### Murine model of cryptococcosis

*C. neoformans* H99, A1-84-18, and A1-38-2 were cultured in YPD for 48 hours at 30°C. Harvested cells were washed three times with sterile PBS, enumerated with a hemocytometer and diluted to 1.9×10^6^ CFU/mL in sterile PBS. CD-1 females (Envigo), 25-30 grams, were inoculated with 3.8×10^5^ CFU (200 µL) by tail-vein injection. For fungal burden, brains were harvested 5 days post inoculation, homogenized in sterile PBS (1 mL) and ten-fold dilutions were plated on YPD. Differences between groups were analyzed by one-way ANOVA followed by Tukey’s multiple comparisons test. For virulence studies, mice were monitored for 21 days following *C. neoformans* inoculation. Percent survival was plotted on a Kaplan-Meier curve and a Log-rank test was used to assess statistical significance of the curves.

### Ethics Statement

The Guide for the Care and Use of Laboratory Animals of the National Research Council was strictly followed for all animal experiments. The animal experiment protocols were approved by Institutional Animal Care and Use Committee at the University of Iowa (protocol: 7102064).

### *Galleria mellonella* larva model of cryptococcosis

The larvae of the great wax-moth *G. mellonella* in the final instar larval stage were obtained from Vanderhorst, Inc. (St. Marys, OH, USA). The *G. mellonella* larvae (0.3–0.4 g) were used for inoculated as previously described (19). Briefly, 1×10^5^ *C. neoformans* cells in PBS (5 µl) were injected into the hemocoel of each wax-moth via the last left proleg. After injection, the wax-moth larvae were incubated at 37°C in dark. For each experiment, 10-15 wax-moth larvae per group were infected and monitored for survival. Kaplan-Meier curves were analyzed using Log-rank test to determine statistical significance for differences between groups.

### E-test and disk-diffusion assay

Briefly, yeast cells at a cell density of approximately 5×10^6^ were spread onto RPMI 1640 agar medium with L-glutamine and without sodium bicarbonate. The plates were allowed to dry. In disk diffusion assays, Whatman paper discs (7 mm) containing DMSO, fluconazole, itraconazole and myriocin at indicated concentrations were dried and placed on the agar surface. E-test strips (bioMérieux) with amphotericin B, fluconazole, or voriconazole were placed on the agar surface. The plates were incubated at 37°C in ambient air or at 5% CO_2_.

## Acknowledgements

We thank Zhiyao Yang for performing Candida albicans susceptibility testing. We gratefully acknowledge the financial support from the Biology department of Texas A&M University, the Microbiology department of the University of Georgia (startup funds to XL), and the Department of Pediatrics (start-up funds for DJK). Dr. Lin holds an Investigator Award in the Pathogenesis of Infectious Disease from the Burroughs Wellcome Fund (<u>1012445</u> to XL).

---

**Supplemental Figure 1. Clinical isolates are generally CO_2_-tolerant.** A set of 10 clinical *C. neoformans* var. *grubii* isolates from Duke University were spotted on RPMI media as described in Fig. 1. The images were taken after 3 days at 37^°^C in either ambient air or 5% CO_2_. The images are representative of 2 biological replicates that showed identical phenotypes.

**Supplemental Figure 2. Glucose supplementation does not confer CO_2_ tolerance to CO_2_ sensitive strains.** Three CO_2_ sensitive environmental strains A2-102-5, A4-34-6, and A5-35-17 were incubated on RPMI medium or RPMI+2% glucose medium at 37°C in ambient air or in 5% CO_2_.

**Supplemental Figure 3. An environmental CO_2_ sensitive strain replicates slowly in brain**. All animals infected with A1-38-2 survived to day 21 and were sacrificed. The brain fungal burden was determined as described in Fig. 1. The data are plotted with those from **Fig. 1D** and indicate an 1.5 log increase in burden over the 16 days between harvest time points.

**Supplemental Figure 4. CO_2_ affects cryptococcal susceptibility to antifungal drugs.** (**A**) H99 is more susceptible to fluconazole in 5% CO_2_ at all concentrations tested by disk diffusion assay. H99 cells were spread onto RPMI agar medium. Disks containing fluconazole (2μg, 4.5μg, 6μg, or 10μg) or DMSO (control) were air dried and placed on top of the yeast lawn. Cells were incubated at 37°C for 2 days in ambient air or in 5% CO_2_. (**B**) The CO_2_-tolerant clinical isolate C23 showed increased susceptibility toward fluconazole (FL) and voriconazole (VO) in CO_2_, but its susceptibility towards amphotericin B remained similar either in ambient air or in CO_2_. A cell suspension of strain C23 was spread onto RPMI agar medium. E-test strips with amphotericin B (AP), fluconazole (FL), or voriconazole (VO) were placed on top of the air-dried yeast lawn. The cells were then incubated at 37°C in ambient air or in 5% CO_2_.

## References

1. Rajasingham R, Smith RM, Park BJ, Jarvis JN, Govender NP, Chiller TM, Denning DW, Loyse A, Boulware DR (2017) Global burden of disease of HIV-associated cryptococcal meningitis: an updated analysis. Lancet Infect Dis. 17:873–881.

2. Maziarz EK, Perfect JR (2017) Cryptococcosis. Infect Dis Clin N Am 30:179–206.

3. May RC, Stone NR, Wiesner DL, Bicanic T, Nielsen K (2016) *Cryptococcus*: from environmental saprophyte to global pathogen. Nat Rev Microbiol 14:106–117.

4. Litvintseva AP, Mitchell TG (2009) Most environmental isolates of *Cryptococcus neoformans* var. *grubii* (serotype A) are not lethal for mice. Infect Immun 77:3188–3195.

5. Mukaremera L, McDonald TR, Nielsen JN, Molenaar CJ, Akampurira A, Schutz C, Taseera K, Muzoora C, Meintjes G, Meya DB, Boulware DR, Nielsen K (2019) The Mouse Inhalation Model of *Cryptococcus neoformans* Infection Recapitulates Strain Virulence in Humans and Shows that Closely Related Strains Can Possess Differential Virulence. Infect Immun 87:pii.e00046–19.

6. Kronstad J, Jung WH, Hu G (2008) Beyond the big three: systematic analysis of virulence factors in *Cryptococcus neoformans*. Cell Host Microbe 4:308–310.

7. Casadevall A, Coelho C, Cordero RJB, Dragotakes Q, Jung E, Vij R, Wear MP (2018) The capsule of *Cryptococcus neoformans*. Virulence 1–10.

8. Granger DL, Perfect JR, Durack DT (1985) Virulence of *Cryptococcus neoformans*: regulation of capsule synthesis by carbon dioxide. J Clin Invest. 76:508–516.

9. Odom A, Muir S, Lim E, Toffaletti DL, Perfect J, Heitman J (1997) Calcineurin is required for virulence of *Cryptococcus neoformans*. EMBO J 16:2576–2589.

10. Bahn YS, Cox GM, Perfect JR, Heitman J (2005) Carbonic anhydrase and CO_2_ sensing during *Cryptococcus neoformans* growth, differentiation, and virulence. Curr Biol 15:2013–2020.

11. Lin X, Litvintseva AP, Nielsen K, Patel S, Floyd A, Mitchell TG, Heitman J (2007) alpha AD alpha hybrids of *Cryptococcus neoformans*: evidence of same-sex mating in nature and hybrid fitness. PLOS Genet 3:1975–1990.

12. Jones RP, Greenfield PF (1982) Effect of carbon dioxide on yeast growth and fermentation. Enzyme Microb Technol 4:210–223.

13. Endeward V, Arias-Hidalgo M, Al-Samir S, Gerolf G (2017) CO_2_ permeability of biological membranes and role of CO_2_ channels. Membranes 7:61.

14. Krysan DJ (2015) Toward improved anti-cryptococcal drugs: novel molecules and repurposed drugs. Fungal Genet Biol. 78:93–98.

15. Grossman NT, Casadevall A (2017) Physiological Differences in *Cryptococcus neoformans* Strains *In Vitro* versus *In Vivo* and Their Effects on Antifungal Susceptibility. Antimicrob Agents Chemother 61: pii: e02108–16.

16. Sherman F (2002) Getting started with yeast. Methods Enzymol. 350:3–41.

17. Upadhya R, Lam WC, Maybruck BT, Donlin MJ, Chang AL, Kayode S, Ormerod KL, Fraser JA, Doering TL, Lodge JK (2017) A fluorogenic *C. neoformans* reporter strain with a robust expression of m-cherry expressed from a safe haven site in the genome. Fungal Genet Biol. 108:13–25.

18. Fan Y, Lin X (2018) Multiple Applications of a Transient CRISPR-Cas9 Coupled with Electroporation (TRACE) System in the *Cryptococcus neoformans* Species Complex. Genetics 208:1357–1372.

19. Mylonakis E, Moreno R, El Khoury JB, Idnurm A, Heitman J, Calderwood SB, Ausubel FM, Diener A (2005) *Galleria mellonella* as a model system to study *Cryptococcus neoformans* pathogenesis. Infect Immun 73:3842–3850.

